# Sensory feedback-dependent coding of arm position in local field potentials of the posterior parietal cortex

**DOI:** 10.1101/2020.09.11.293365

**Authors:** Paul VanGilder, Ying Shi, Gregory Apker, Christopher A. Buneo

## Abstract

Although multisensory integration is crucial for sensorimotor function, it is still unclear how sensory cues provided by the visual and proprioceptive systems are combined in the brain during motor behaviors. Here we characterized the effects of multisensory interactions on local field potential (LFP) activity obtained from the superior parietal lobule (SPL) as non-human primates performed an arm position maintenance task with either unimodal (proprioceptive) or bimodal (visual-proprioceptive) sensory feedback. Based on previous analyses of spiking activity, and observations that LFP and spikes are highly correlated in some cortical areas, we hypothesized that evoked LFP responses would be tuned to arm location but would be suppressed on bimodal trials, relative to unimodal trials. We also expected to see a substantial number of recording sites with enhanced beta band spectral power for only one set of feedback conditions, as was previously observed for spiking activity. We found that evoked activity and beta band power were tuned to arm location at many individual sites, though this tuning often differed between unimodal and bimodal trials. At the population level, both evoked and beta band activity were consistent with feedback-dependent tuning to arm location, while beta band activity also showed evidence of suppression on bimodal trials. The results suggest that multisensory interactions can alter the tuning and gain of arm position-related LFP activity in the SPL and that this activity can be used to decode the arm’s location under varying sensory conditions.

## Introduction

Multisensory (or multimodal) integration (MSI) is crucial for sensorimotor function, particularly when estimating the state of the body (i.e. the position and velocity of relevant body segments) and planning movements. Combining multiple sensory cues provides a means of overcoming inherent noise in the sensory systems and reduces uncertainty in state estimates (Ernst and Bülthoff, 2004). This has been demonstrated across a variety of behavioral domains including target localization, object recognition, navigation, and limb movement (Giard and Peronnet, 1999; Ernst and Banks, 2002; Battaglia et al., 2003; Sober and Sabes, 2003; Bremmer, 2005; Morgan et al., 2008). For example, during reaching movements, visual and proprioceptive cues are thought to be combined with hand position and velocity estimates derived from efference copies of motor commands and a forward model to form estimates of hand location that are less biased and more precise (Wolpert et al., 1995; Van Beers et al., 1996).

However, it is still unclear how sensory cues are combined in the brain. Seminal neurophysiological studies of subcortical neurons by Stein and colleagues quantified the degree of enhancement or suppression of neuronal firing rates arising during MSI (Stein and Meredith, 1993; Wallace and Stein, 1997; Jiang et al., 2001). These studies helped establish the well-known principles of inverse effectiveness (i.e., greater enhancement for weaker stimuli) and superadditivity (i.e. multisensory responses greater than the sum of unimodal responses) (for review see: Stein and Stanford, 2008). Although these early studies emphasized responses to weak unimodal stimuli, and therefore multisensory enhancement of spiking activity, more recent investigations of the cerebral cortex suggest subadditivity (including suppression) may be more common (Sugihara et al., 2006; Avillac et al., 2007; Morgan et al., 2008; Kayser et al., 2010; Shi et al., 2013). Importantly, suppression of activity under bimodal conditions has been shown to be associated with greater information transmission (Kayser et al., 2010) and improved decoding accuracy (Shi et al., 2013). Moreover, a network level divisive normalization model has been developed that can account for the empirical principle of inverse effectiveness and observed sub- and superadditivity (Oshiro et al. 2011) and probabilistic population codes predict subadditive (suppressed) neural responses during Bayesian integration of sensory cues (Ma et al., 2006).

More recent work indicates that MSI may manifest in the brain through other mechanisms as well. For example, changes in neuronal spike timing or variability have been observed during multisensory interactions (Kayser et al., 2010; Koehler et al., 2011; Shi et al., 2013; Chabrol et al., 2015; VanGilder et al., 2016) and local field potentials (LFPs) have been shown to exhibit both multisensory enhancement and suppression. For example, in non-human primates, LFPs are either enhanced or suppressed depending on the temporal congruency and type of facial and vocal expressions of conspecifics (Ghazanfar, 2005). Similarly, enhancement or suppression of LFPs can occur depending on whether audio-visual cues are derived from conspecifics, other animals, or artificial sources. (Kayser et al., 2008). Modulations of spectral power within specific frequency bands of EEGs and LFPs during multisensory processing have also been observed (Belitski et al., 2010; Sarko et al., 2013; Engel et al., 2007, 2012). Regarding visual-proprioceptive interactions specifically, although effects of these interactions on spike rates and spike timing have been previously described (Graziano et al., 2000; Shi et al., 2013; VanGilder et al., 2016), little information exists regarding the effects of such multisensory interactions on LFPs.

We have previously characterized multisensory interactions in a population of neurons in the superior parietal lobule (SPL) as non-human primates performed an arm position maintenance task with unimodal (proprioceptive) or bimodal (visual-proprioceptive) sensory feedback (Shi et al., 2013; VanGilder et al., 2016). Our rationale for focusing on activity related to position maintenance rather than movement was that multisensory interactions should be more amenable to study under quasi-static rather than highly dynamic conditions. Moreover, study of the neural substrates and mechanisms of arm position maintenance have been relatively ignored, despite evidence that they may only partially overlap with those involved in movement (Brovelli et al., 2005; Shadmehr, 2017). Regarding effects of multisensory interactions, we found that, relative to unimodal conditions, neuronal firing rates were largely suppressed under bimodal conditions, consistent with subadditive multisensory interactions. Some neurons also exhibited beta (13-30Hz) oscillatory spiking under only one set of sensory conditions (unimodal or bimodal), while others did so under both conditions. In the current study, we examined LFPs recorded during the same experiments. For our previous analyses of spike times, we focused on the beta band due to the prevalence of strong oscillatory activity in this range in posterior parietal areas during various tasks (Buneo et al., 2003; Joelving et al., 2007; Witham and Baker, 2007) and its reported role in linking large-scale cortical networks during the maintenance of sensorimotor state (Brovelli et al., 2004; Engel and Fries, 2010). We hypothesized that patterns of enhancement and suppression observed in spiking activity would be reflected in evoked LFP responses, i.e. these responses would be suppressed on bimodal trials relative to unimodal trials. In addition, we expected to see modulations of LFP power in the beta band that mirrored those observed in the spike spectra, i.e. beta power would be enhanced at individual recording sites under one or both sets of conditions.

## Methods

### Experimental Subjects and Paradigm

All experimental and veterinary care procedures were approved by the Arizona State University Institutional Animal Care and Use Committee and conducted according to the U.S. Public Health Service Policy on Humane Care and Use of Laboratory Animals (Public Law 99–158) and the Guide for the Care and Use of Laboratory Animals (National Academy Press, 1996). Environmental social enrichment, housing, and feeding procedures also conformed to institutional standards, which are AAALAC International accredited.

We have described the experimental paradigm and apparatus in previous reports but provide an overview here (Shi et al., 2013; VanGilder et al., 2016). Briefly, two rhesus macaques (‘X’ and ‘B’) were trained to make reaching movements within a semi-immersive 3D virtual reality environment displayed on a 3D monitor and projected onto a mirror in their fields of vision. The monkeys made center-out reaches to eight peripheral targets and maintained their hand location at these targets with or without visual feedback. The mirror blocked the view of each monkey’s actual arm, but visual feedback of hand location was provided as a spherical cursor within the virtual environment. An active motion tracking system (Phoenix Technologies Inc.) monitored arm movements via LED markers placed on each monkey’s wrist. Eye movements were tracked using a remote optical tracking system (Applied Science Laboratories, Inc.). At the start of each trial, an animal had to align the arm cursor with the starting location, which appeared as a green sphere presented in the center of the virtual workspace. Once this location had been maintained for 500ms (baseline period), one of the peripheral reach targets was pseudorandomly presented, serving as the “go” cue to begin the reach (movement period). When the peripheral target was acquired, an animal performed a saccade back to the starting location. This began the “static holding period,” where the animal maintained its hand location at the peripheral target while fixating at the starting location for 800-1200ms. During the static holding period, visual feedback of the arm cursor was allowed on half the trials (bimodal (B) condition) and removed on the remainder (unimodal (U) condition). Spherical behavioral windows with radii ranging from 2-2.4 cm surrounded the reach targets and a behavioral window with a radius of ~6.5° of visual angle surrounded the fixation point. Trials were deemed successful if the animals acquired both the reach targets and fixation point and maintained position within these windows for the remainder of the trial. Animals completed five (5) successful trials to each target in both sensory conditions and target locations were pseudorandomly varied on a trial-by-trial basis.

### Data Acquisition

We analyzed evoked LFP responses from 173 recording sites (97 from monkey X, and 73 from monkey B) located within the superficial cortex of the superior parietal lobule (area 5). Note that this is less than the number of recording sites reported in Shi et al. (2013) and VanGilder et al. (2016) due to technical issues that corrupted the signals at some sites. LFPs were recorded acutely using varnish-coated tungsten microelectrodes (~1-2MΩ at 1kHz). LFPs were separated from the spike data after amplification by low-pass filtering at 300Hz and were sampled at 1kHz before saving to disk with the associated behavioral data (Multichannel Acquisition Processor, Plexon Inc.).

### Data Analysis

All analyses were conducted in MATLAB (MathWorks, Natick, MA). For all statistical analyses, an alpha of 0.05 was used.

#### Evoked Potentials

LFP data underwent two stages of preprocessing. The Chronux toolbox (Bokil et al., 2010, http://chronux.org) was used to remove line noise and slow voltage fluctuations caused by electrical transients that may cause a slow “drift” of the signal. Line noise was removed using Thomson’s regression method to detect and remove 60Hz sinusoids and any harmonics from the data (Jarvis and Mitra, 2001; Bokil et al., 2010). A sliding-window linear regression procedure was used to remove the slow voltage drift wherein a least-squares trend line was fit to the signal within each successive temporal window. Subsequently, the best fitting trend lines in each window were then weighted and combined to estimate the slow fluctuation, which was then removed from the data signal.

To examine the effects of hand location and sensory condition on evoked responses, the filtered LFP signals were first aligned to the start of the holding period. For each hand location, the mean LFP response for each trial and sensory condition was squared, and then averaged over the holding period time window. The square root of this quantity (RMS) was then compared across final hand locations and sensory conditions (Liu and Newsome, 2006; O’Leary and Hatsopoulos, 2006). Specifically, a two-way ANOVA (factors: hand location, sensory condition) was used to assess the effects of sensory condition and reach direction on the mean evoked LFP response at individual recording sites during the baseline, movement, and holding periods.

Using the same framework from previous experiments (Stein et al., 1989; Shi et al., 2013) an enhancement/suppression index was also computed for the evoked LFP responses obtained at each site. First, a preferred hand location was determined. Following convention used in previous studies, the preferred location was defined as the hand location with the largest trial-averaged evoked response in the U condition (O’Leary and Hatsopoulos, 2006; Shi et al., 2013). Next, LFP responses were averaged across trials for the preferred hand location for both B and U conditions. Enhancement/suppression indexes (I) were computed as follows:

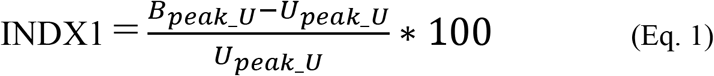

where B and U refer to the trial-averaged evoked responses for the preferred hand locations (as defined by the U condition). For this index, positive values indicate enhancement and negative values indicate suppression of LFP responses under bimodal conditions. Suppression or enhancement was considered to be statistically significant based on the result of the ANOVA performed on the RMS values. We also calculated a second index (INDX2) to account for the possibility that some enhancement/suppression might arise from differences in tuning between the two conditions:

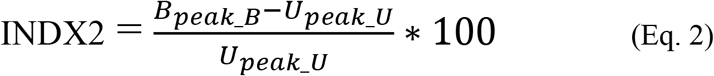

We also analyzed population-level differences in the evoked responses. For this, evoked responses were first normalized by baseline activity then averaged across trials and recording sites for the preferred location in both conditions. For these analyses, a nonpreferred location was also determined and was defined as the hand location with the lowest trial-averaged evoked response in the U condition. T-tests were used to determine if the evoked responses differed between B and U conditions, and between preferred and nonpreferred locations for each condition.

#### Spectral Analysis

Temporal structure in the LFPs was analyzed using the multitaper spectral estimation technique. For each trial, we used 9 data tapers and a time-bandwidth parameter of 5, providing a spectral resolution of 6.25Hz. For each recording site, trial-averaged power spectra with jackknife error bars were computed for each reach direction/hand location and condition. At the population level, spectra for each recording site were normalized by the average baseline power (−500-0 ms prior to target onset) before averaging across sites. Population spectra error bars were derived from the jackknife standard error across recording sites. All analyses were performed using custom MATLAB programs supplemented by the Chronux toolbox (Bokil et al., 2010) for multitaper spectral analyses.

We also analyzed the average LFP spectral power within each of the following frequency bands: delta (0-4Hz), theta (4-8 Hz), alpha (8-12 Hz), beta (13-30 Hz), and gamma (30-90 Hz). As with the evoked responses, for each recording site a two-way ANOVA (reach direction, sensory condition) was conducted to assess the effects of reach direction/hand location and sensory condition on the power in a given band. At the population level, T-tests were used to test for differences between population spectra associated with each sensory condition, as well as between preferred/nonpreferred locations for each condition.

## Results

### Behavior

Analyses of behavioral data were previously reported (Shi et al., 2013) but will be summarized here. The experimental paradigm maximized the likelihood that final arm locations would be the same during both B and U conditions of the task. This was to ensure that any observed changes in neural responses could be interpreted as resulting from interactions between sensory cues and not differences in endpoint locations. Mean endpoint locations and variances along the horizontal, vertical and depth axes did not differ significantly between sensory conditions, suggesting that during the static holding period, hand location was largely identical in the two conditions. The results of the current study were interpreted within this context.

### Time Domain

Results of a two-way ANOVA (factors: sensory condition and hand location/movement direction) on the mean evoked LFP responses recorded at individual sites during the baseline, movement, and holding periods are summarized in Table 1.

**Table 1.**
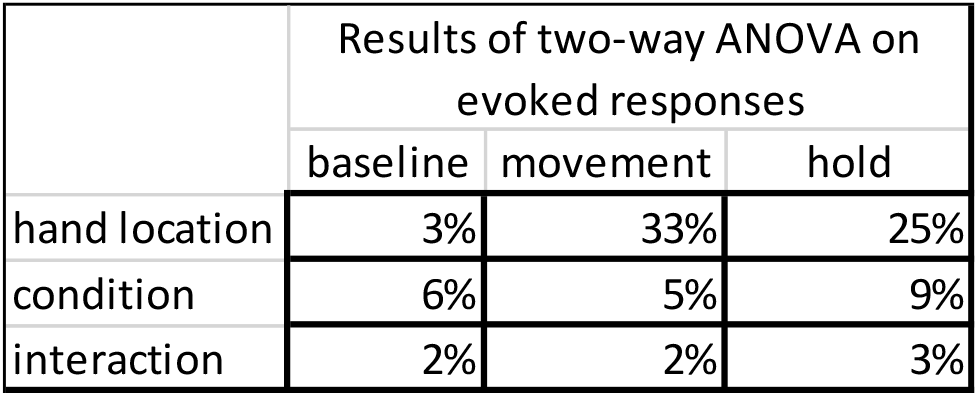
Summary of two-way ANOVA on baseline, movement, and hold epochs. Numbers represent percentages of recording sites (n=173) that showed significant effects (p<0.05).

Main effects of hand location were common during the 800-ms static holding period, though at a smaller number of sites (43/173; ~25%) than during the movement period (57/173; ~33%). Figure 1 shows the LFP evoked responses at an exemplary recording site that demonstrated significant effects of hand location during the holding period. The voltage traces for both sensory conditions were largely similar at each hand location, though there were clear differences among the responses at the eight target locations. Looking at the central polar plot, mean LFP evoked potentials were greater for hand locations up and to the left of the starting location, with a maximal response at 135°.

**Figure 1.**
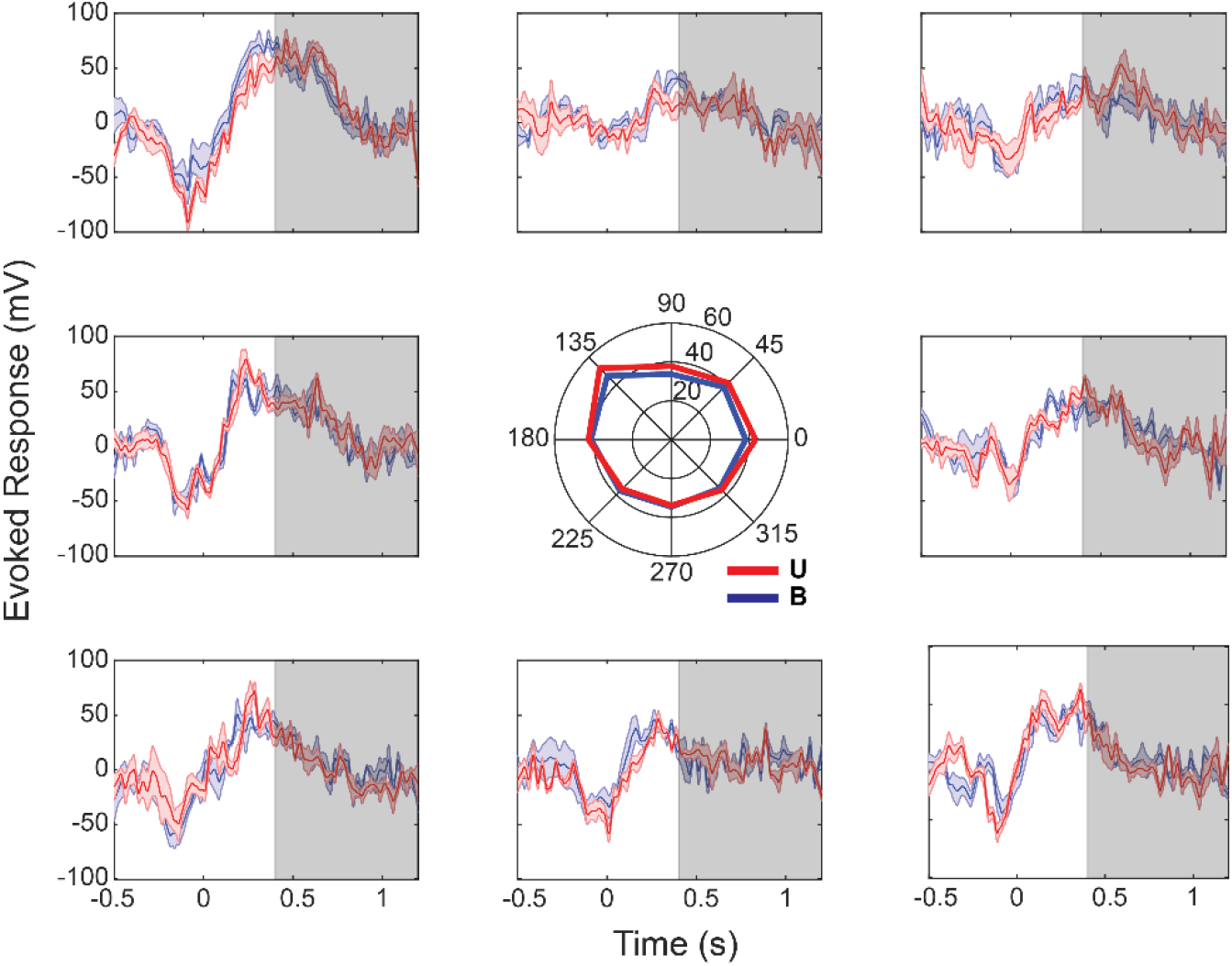
Evoked LFP responses at a recording site with only significant effects of hand location during the holding period. Each panel corresponds to one of the 8 reach targets. Trial averaged responses (with bootstrap error bars) aligned to reach target acquire are shown. Grey box corresponds to the static holding period (0.4-1.2s) after target acquire.

Although only a relatively small percentage of individual recording sites (~9%) exhibited statistically significant effects of sensory condition during the holding period by ANOVA, an analysis of multisensory enhancement/suppression indexes (INDX1; Eq. 1) calculated at each recording site revealed that activity during B trials was generally suppressed relative to activity on U trials.**Error! Reference source not found**. Figure 2 shows a bar graph of these indices for all recording sites. For the preferred hand location, over 87% had a negative index value (mean index value= −20.6), indicating activity was largely suppressed during B trials relative to U trials. Importantly, INDX1 reflects differences in activity between conditions at the preferred location defined by the *U* condition. Thus, this index assumes that tuning was identical in the two conditions. If this was not the case, then some of the observed suppression could reflect differences in spatial tuning to hand location rather than differences in the sensory conditions themselves. When tuning was compared between conditions at individual sites, differences were common: 123 recording sites (71%) had different tuning for the two sensory conditions. Since reaches were directed to 8 discrete targets locations, tuning differences between conditions were quantified as relative distance between preferred locations, with a mean of 2.14 targets locations.

**Figure 2.**
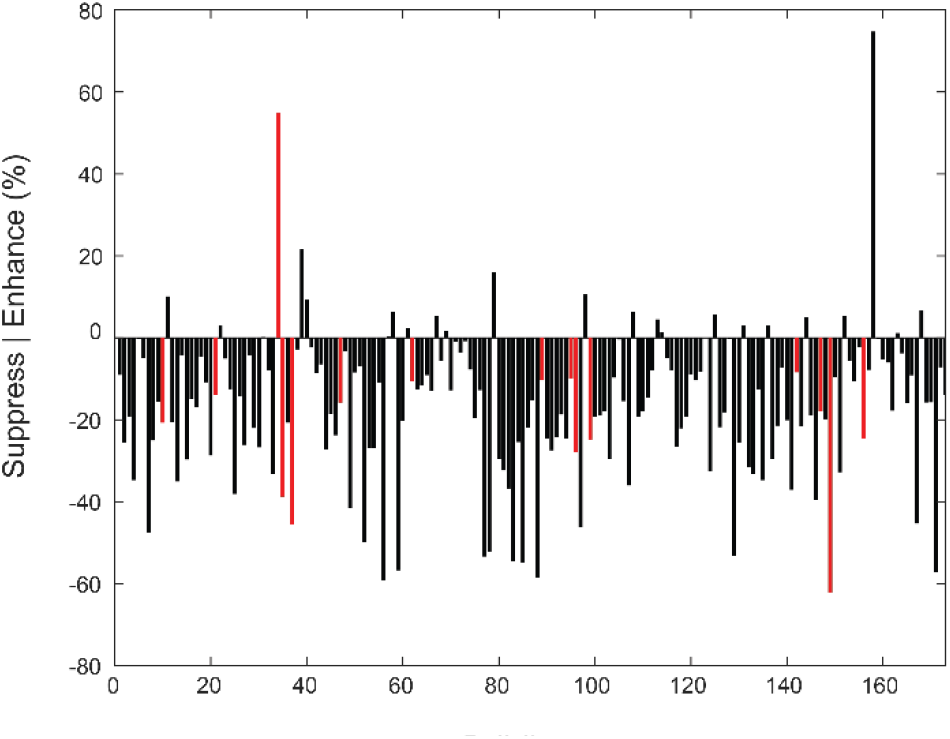
Multisensory interaction indices (INDX1) for evoked responses during the holding period. Red bars indicate recording sites that exhibited statistically significant effects of sensory condition (ANOVA, p<0.05).

To account for the effects of tuning changes on multisensory enhancement/suppression, we also calculated a second index that was based on the preferred location in *each* sensory condition (INDX2). Here, suppression was not as common nor as strong (mean index value = −13.8), though activity was still suppressed at a majority of recording sites during the B condition (not shown). Nevertheless, this suggests that many individual sites demonstrated differences in activity between conditions that could have reflected a combination of tuning differences and multisensory response suppression or just tuning differences alone.

Tuning differences, multisensory suppression or a combination of the two effects lead to different predictions regarding population level analyses focused on the preferred direction. For example, Figure 3 shows idealized tuning curves for hand location for both conditions (Georgopoulos et al., 1984; Todorov, 2002; Lalazar et al., 2016). Four scenarios are illustrated. In Figure 3A, the curves exhibit tuning differences between conditions without attenuation of responses in the B condition, i.e. without multisensory suppression. The bar plots to the right show the expected differences in activity when the preferred location for the U condition (PL_U_) is used to compare responses as well as when the preferred location for the B condition (PL_B_) is used. Under this scenario, differences in activity at the preferred location are expected to be equivalent in magnitude but opposite in sign for the two comparisons. For the scenario illustrated in Figure 3B, tuning is assumed to be identical for the two conditions but with suppression of response magnitude in the B condition. Here, differences in response magnitudes are expected to be equivalent in magnitude *and* sign for comparisons based on both PL_U_ and PL_B_. In **Figure 3 Error! Reference source not found**.C and Figure 3D, combinations of tuning differences and bimodal suppression are shown. In these scenarios, differences in response magnitudes for the two comparisons are nonequivalent in magnitude with signs that depend on the difference in tuning. For small differences in tuning, signs are expected to be the same (Figure 3C), while for larger differences they are expected to differ (Figure 3D). In conclusion, condition-dependent differences in activity can be distinguished from differences in tuning by comparing the magnitude and sign of response differences between PL_U_ sorted and PL_B_ sorted datasets, a strategy that was employed in the analysis of our population evoked potentials and spectra.

**Figure 3.**
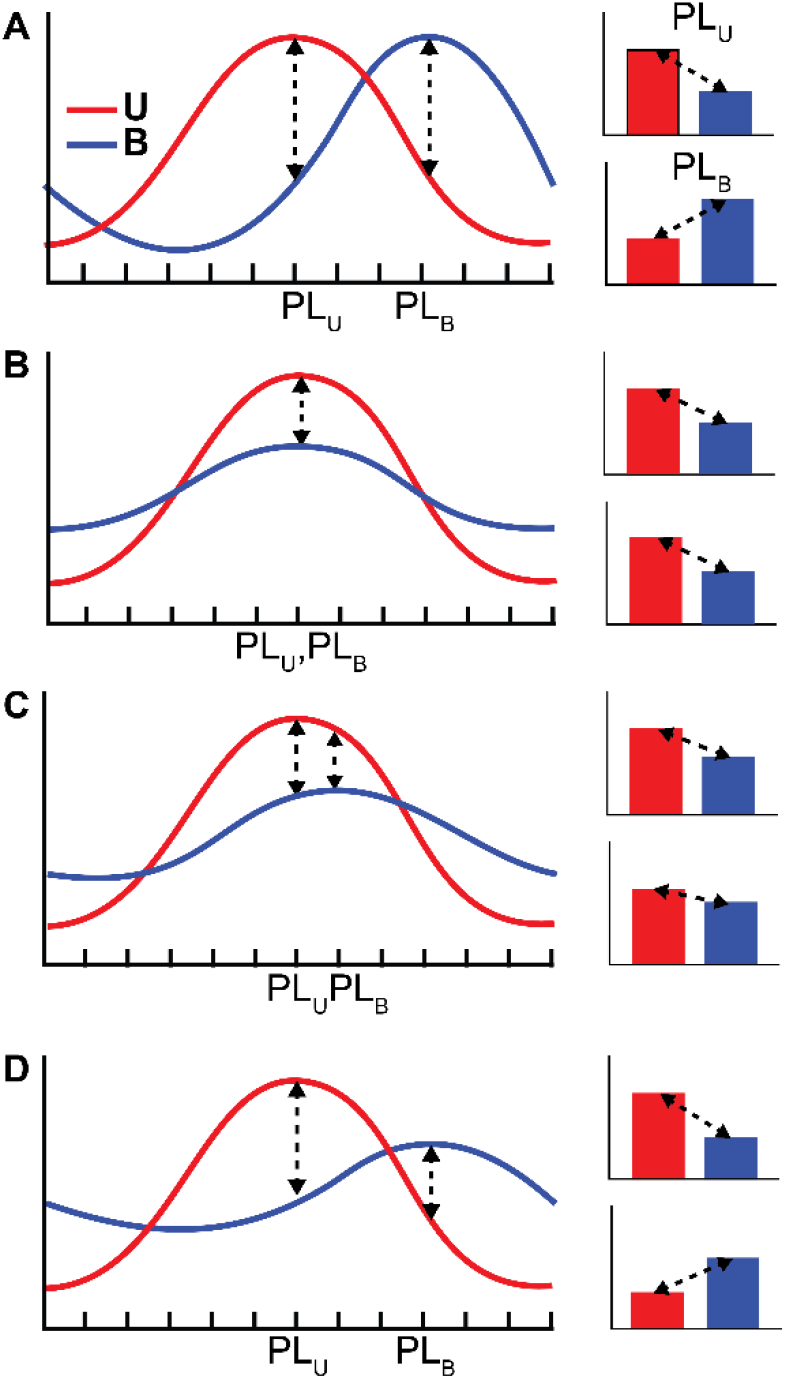
Idealized cosine-tuning curves for hand location in the presence and absence of multisensory suppression and/or tuning changes. Cosine tuning functions were assumed based on (Georgopoulos et al., 1984) (see also (Lalazar et al., 2016). A: Shift in tuning with no suppression. B: Suppression without tuning changes. C: Suppression combined with small tuning changes. D: Suppression combined with large tuning changes. See text for details.

Figure 4 shows the population evoked responses for PL_U_ sorted (A) and PL_B_ (B) sorted datasets. Activity for both sensory conditions and hand locations (preferred and nonpreferred) are shown. Generally speaking, LFP activity prior to movement onset (~ −0.4s) was indistinguishable between sensory conditions and hand locations. During the subsequent movement period, the temporal profiles for each sensory condition were largely similar for their respective reach directions, though response magnitudes differed between directions (as expected). Although responses were sorted into preferred or nonpreferred based on activity during the *holding* period, these earlier differences in magnitude indicate that the evoked LFPs were also strongly modulated by movement direction (see also Table 1). Notably though, no differences between sensory conditions were observed prior to or during movement.

**Figure 4.**
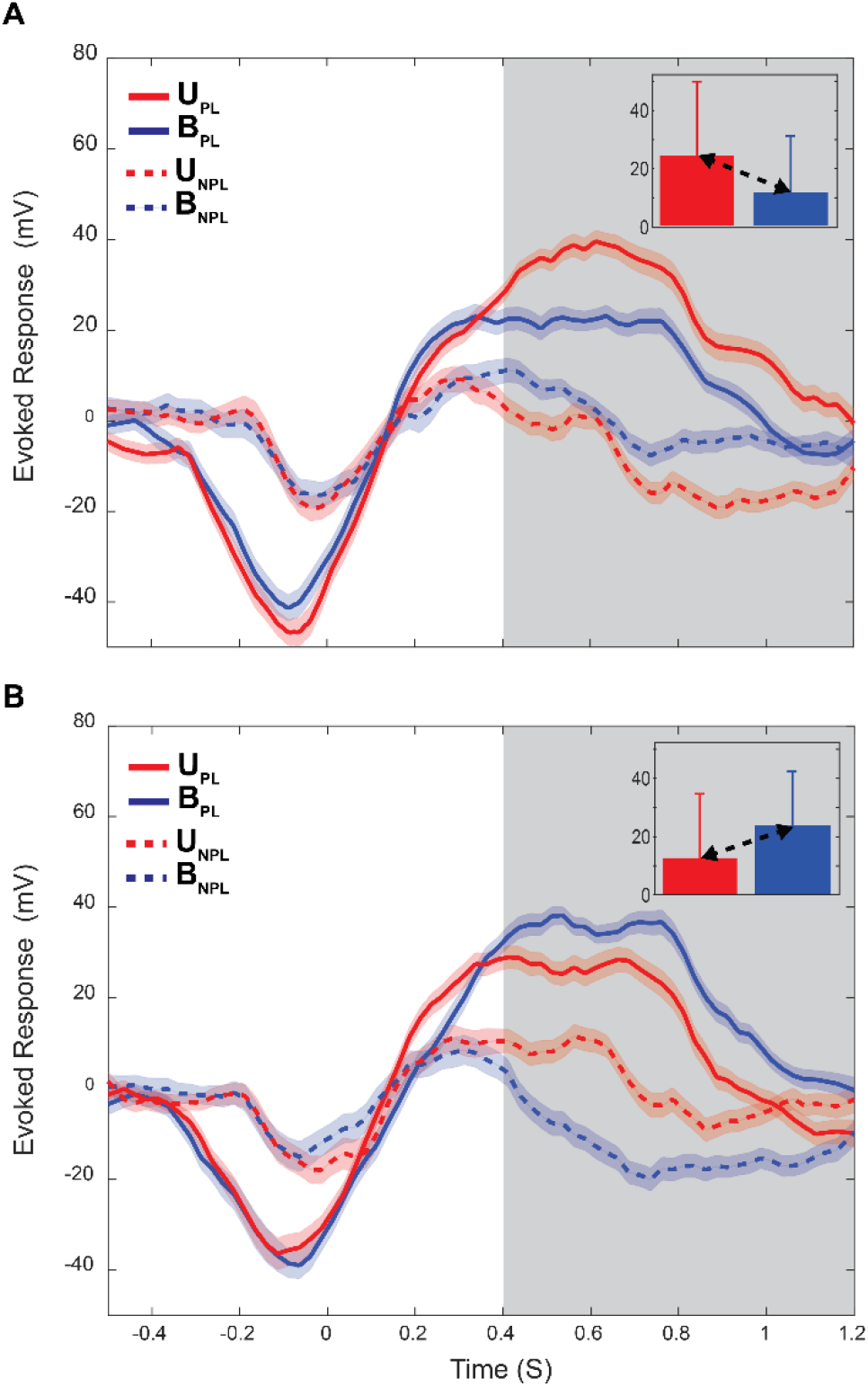
Mean population evoked responses for both sensory conditions and hand locations (preferred and non-preferred). Data are aligned to target acquire (t=0). The grey box (04.-1.2s) corresponds to the 800-ms static holding period. A: Responses when the U condition was used to define preferred location. Inset: means and standard deviations over the entire holding period. B: Responses when the B condition was used.

Near the start of the static holding period (Fig. 4, grey boxes) differences in evoked responses due to movement direction *and* sensory condition became apparent. The divergence of activity between the U and B conditions at the start of the holding period is notable because it is the only part of the task where the sensory feedback conditions differed, i.e. visual information about arm location was only available until the start of the holding period in the U condition. Previous analyses established that the distributions of hand locations were largely similar between conditions during this period of the task (Shi et al., 2013), thus the differences in LFP activity seen here could be attributed to differences in sensory condition, differences in tuning, or both (as previously discussed). For the PL_U_ sorted data (Fig. 4A), evoked activity for the preferred hand location differed significantly between U and B conditions during the holding period (*t* test, p<0.05), with activity for the B condition appearing suppressed with respect to the U condition. For the nonpreferred hand location, activity also differed between conditions during the holding period (*t* test, p<0.05) with activity during the B condition appearing greater (less negative) than that in the U condition.

For the PL_B_ sorted data (Fig. 4B), the opposite trends were observed. That is, although activity also differed significantly between the U and B conditions for both the preferred and non-preferred locations (*t* tests, p<0.05), activity in the B condition appeared *greater* than activity in the U condition for the preferred location and was more negative in the nonpreferred location. When the unsigned differences in activity between conditions were compared statistically between PL_U_ sorted and PL_B_ sorted datasets, no significant differences were found (*t* tests, p<0.05). Thus, at the population level, evoked responses were largely consistent with scenario A in Fig. 3, indicating differences in activity between conditions reflected mainly differences in tuning and not multisensory suppression.

### Frequency Domain

Previous results from this area showed that during the maintenance of static arm positions, the spike trains of many neurons were strongly oscillatory in the beta band (13-30Hz), though modulation of spike timing within other frequency bands was observed as well (VanGilder et al., 2016). As a result, in the present study we analyzed the delta, theta, alpha, beta, and gamma bands of the LFP during the holding period. Figure 5 shows example single site LFP power spectra for the preferred hand location. Spectra obtained during the baseline period (black) are superimposed on spectra obtained during the holding period for both sensory conditions. The shapes of the holding period spectra seen in this figure were typical, with the greatest power being concentrated at the lower frequencies and with a notable ‘bump’ in the beta band. Importantly, power in the lower frequency bands (delta, theta, alpha, beta) generally increased during the holding period relative to that in the baseline period, regardless of sensory condition or final hand location. Nevertheless, power in these frequency bands was modulated by sensory condition and/or hand location at some sites, as described below.

**Figure 5.**
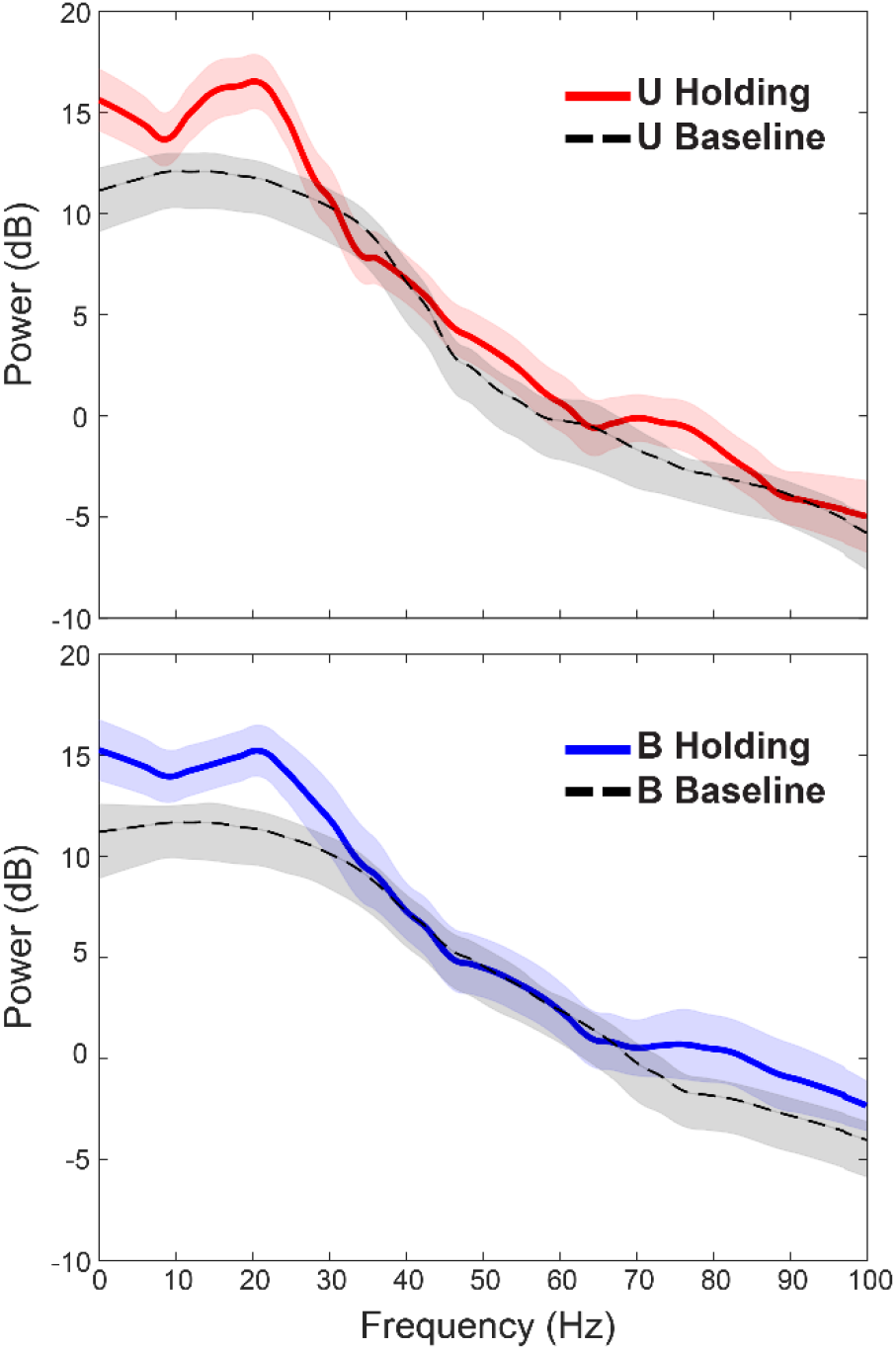
Trial averaged LFP power spectra (with jackknife error bars) for the holding and baseline periods at a single site. Power was enhanced relative to baseline during the holding period from frequencies less than ~30 Hz.

As with the evoked potentials, a two-way ANOVA was used to quantify the effects of sensory condition and final hand location at individual recording sites. Table 2 summarizes the ANOVA results for each frequency band and task condition. Responses across recording sites were largely similar between sensory conditions and hand locations, with many sites having no main effects of either factor, especially in the delta, theta, and alpha bands. Hand location generally had a greater influence on spectral power than did sensory condition during the holding period, particularly at higher frequencies (beta and gamma), where approximately 20% of the sites showed statistically significant effects. The numbers of sites with statistically significant effects of sensory condition were more evenly distributed across frequencies.

**Table 2.** Summary of two-way ANOVA on baseline, movement, and hold epochs. Numbers represent percentages of recording sites (n=173) that showed significant effects (p<0.05).

|  | Results of two-way ANOVA on LFP spectra |  |  |  |  |
| --- | --- | --- | --- | --- | --- |
|  | delta | theta | alpha | beta | gamma |
| hand location | 8% | 6% | 12% | 21% | 24% |
| condition | 6% | 6% | 6% | 6% | 5% |
| interaction | 3% | 2% | 2% | 5% | 3% |

Figure 6 shows the power spectra for an example recording site that showed main effects of hand location in the beta and gamma bands. At this site, beta power was tuned for hand locations down and to the left of the starting position, i.e. for movements that were directed to targets at 225°. However, no observable differences in power between sensory conditions were observed for this frequency band. This was confirmed by ANOVA, which showed a statistically significant effect of hand location (*F*=9.46, *p*<0.05), but no main effect of sensory condition (*F*=0.21, *p*=0.65) nor interaction effects (*F*=0.29, *p*<0.96). Interestingly, at this site power in the lower frequencies was not tuned to direction but tended instead to show effects of the sensory conditions. For example, power in the alpha band was not tuned to final hand location (*F*=1.02, *p*=0.43) but did differ significantly between sensory conditions (*F*=5.85, *p*<0.05), being stronger for the U condition. No significant interaction effect was found (*F*=0.59, *p*=0.77). Thus, when effects of the sensory conditions were observed at single sites they were not necessarily coupled to effects of location/direction but could independent of such effects.

**Figure 6.**
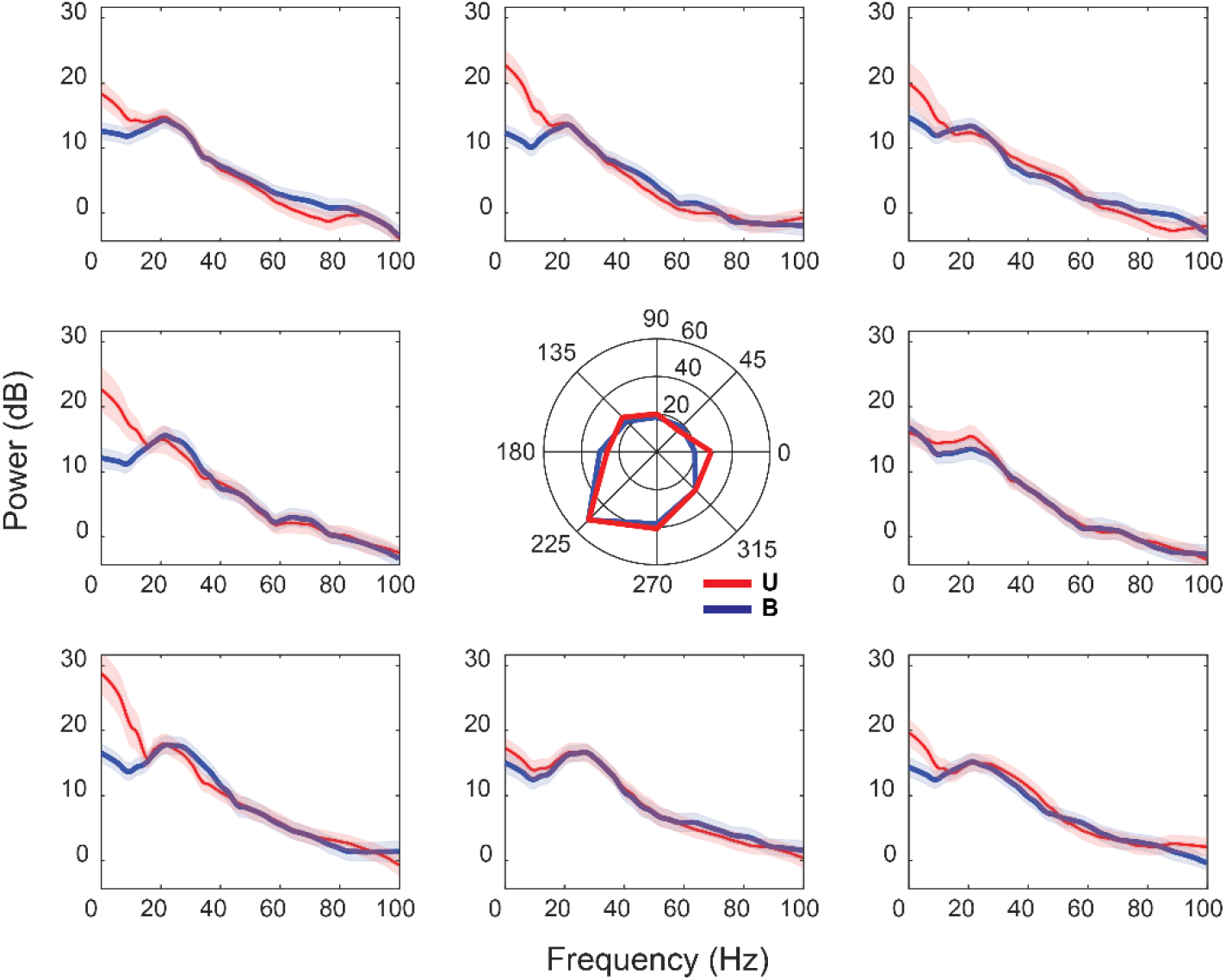
Trial-averaged LFP spectra (with jackknife error bars) from a single recording site that showed main effects of hand location in the beta and gamma bands during the holding period. Spectra were averaged across trials. Blue traces are B trials, red traces are U trials. Spectral power is in log units. Central polar plot is based on mean power in beta band.

As with the evoked activity, even though only a relatively small percentage of individual recording sites (6%) exhibited statistically significant effects of sensory condition on beta power during the holding period, multisensory enhancement/suppression indexes (INDX1) for the preferred hand location indicated that beta band power was generally suppressed on B trials relative to power on U trials. Figure 7 shows a bar graph of these indices for all recording sites. Over 86% of the sites had a negative index value (median index value= −20.45), indicating that beta power was generally greater on U trials than B trials. To assess the extent to which tuning differences between conditions might have played a role in this apparent suppression, we also calculated multisensory enhancement/suppression indices using Eq. 2 (INDX2). As with the evoked responses we found that when using this index suppression was not as common nor as strong (median index value = – 3.45), suggesting that differences in beta power at many sites reflected either a combination of tuning differences and multisensory suppression or tuning differences alone.

**Figure 7.**
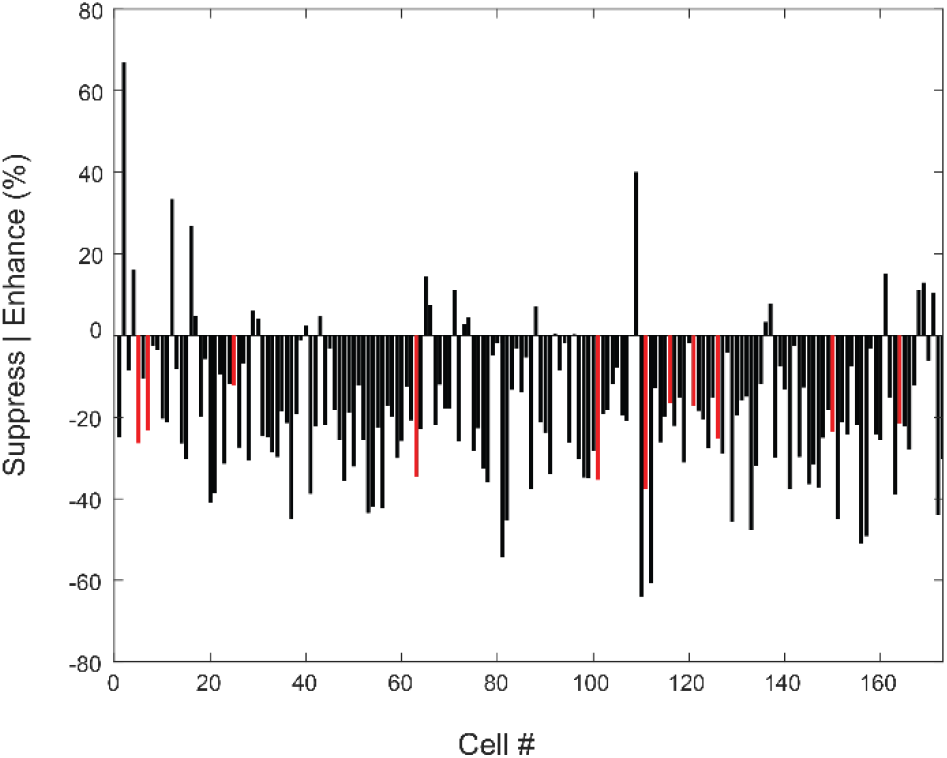
Multisensory interaction indices (Eq. 1) for beta power during the holding period. Red bars indicate recording sites that exhibited statistically significant effects of sensory condition (ANOVA, p<0.05).

Figure 8 shows the population averaged spectra during the holding period for PL_U_ sorted (Figure 8A) and PL_B_ sorted (Figure 8B) datasets. Activity for both sensory conditions and the preferred hand location (based on beta power) are shown. Spectra were normalized by baseline power before averaging. As observed previously, power during the holding period was largely concentrated in the lower frequency bands, with the greatest power being observed in the delta range of frequencies. Power in the theta and alpha frequencies dropped sharply before rising again during the beta band – consistent with the beta bump seen in the single site spectra (Figure 5 & Figure 6).

**Figure 8.**
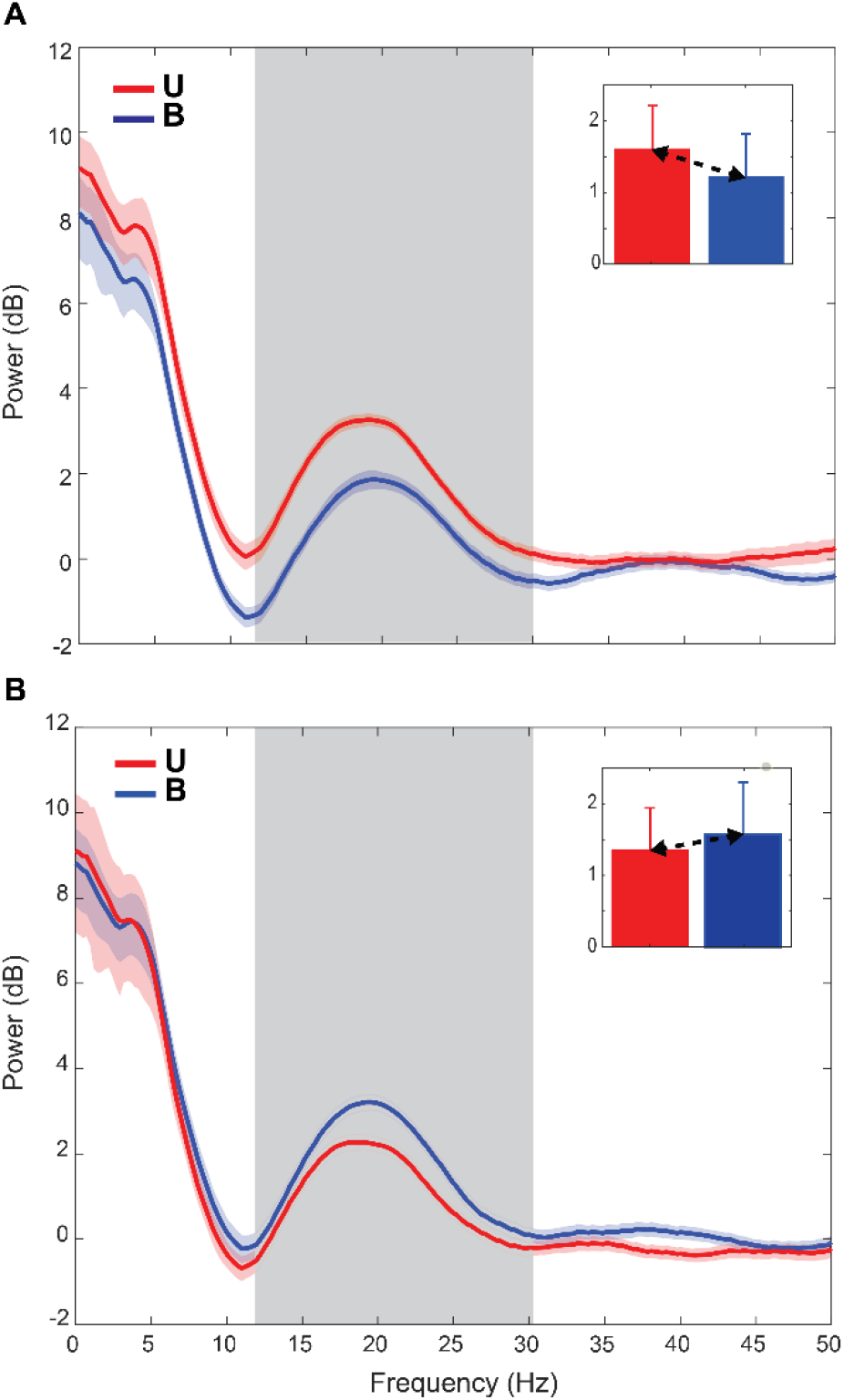
Population spectra for all recording sites in the B (blue) and U (red) conditions during the holding period for the preferred direction. Error bars are jackknife error bars. The grey box indicates the beta band (13-30 Hz). A: Spectra when the U condition was used to define preferred location. Inset: means and standard deviations of the power over the beta band. B: Spectra when the B condition was used.

For the PL_U_ sorted data (Figure 8A), power in the beta band differed significantly between U and B conditions during the holding period (*t* test, p<0.05), with activity for the B condition appearing suppressed with respect to the U condition. Power also differed significantly between the U and B conditions for the PL_B_ sorted data (Figure 8B; *t* test, p<0.05), but in this case power in the B condition was greater than activity in the U condition. Critically, and in contrast to the evoked potentials, unsigned differences in beta power between conditions *differed* between the PL_U_ and PL_B_ sorted datasets (*t* tests, p<0.05). Thus, at the population level, beta power was largely consistent with scenario *D* in Fig. 3, indicating that differences in power between conditions reflected both differences in tuning *and* an overall attenuation of responses in the B condition (multisensory suppression).

## Discussion

Here we examined the effects of bimodal (visual-proprioceptive) interactions on LFP signals recorded at multiple sites in the SPL as non-human primates performed an arm position maintenance task. Based on previous work from our lab and observations that LFPs and spikes are highly correlated in several cortical areas (Pesaran et al., 2008; Denker et al., 2011; Banerjee et al., 2012; Esghaei et al., 2017), we hypothesized that evoked LFP responses would be modulated by multisensory interactions in a manner similar to spiking activity (Shi et al., 2013). That is, we expected that evoked responses would be attenuated on bimodal trials (B) relative to those on unimodal (U) trials. We also expected to see modulations of LFP spectral power that mirrored those observed for spiking activity (VanGilder et al., 2016), i.e. for individual sites beta power was expected to increase during arm position maintenance but for only one set of feedback conditions.

We found that the effects of multisensory interactions on LFP activity (evoked and beta band) were dependent in part upon the criteria used to define preferred hand location. When activity on U trials was used to define the preferred location, multisensory interactions appeared to result predominantly in suppression. However, when activity for both types of trials was used, suppressive effects weakened or reversed sign at many individual sites. At the population level, differences in evoked activity between conditions appeared to result largely from feedback-dependent tuning to hand location. However, for beta band power, effects were more consistent with multisensory suppression superimposed upon feedback-dependent tuning differences. This suggests that different aspects of arm position maintenance (spatial location vs sensory feedback signals available for position) are coupled within the frequency domain representations of LFP activity in the SPL and can be potentially be decoded to extract the current location of the limb, regardless of sensory conditions.

### Multisensory suppression

In the time-domain, evoked activity was suppressed on B trials in a manner similar to spiking activity (Shi et al., 2013), both in degree and in the extent to which tuning differences contributed to this apparent suppression. Previous work has shown that LFP activity and spiking activity are highly correlated in several cortical areas (Siegel et al., 2009; Whittingstall and Logothetis, 2009; Ray and Maunsell, 2011; Khodagholy et al., 2015), though the precise nature of the relationship between these signals remains controversial (Ray, 2015; Pesaran et al., 2018; Watson et al., 2018). For example, although many studies have assumed that LFPs and spikes represent separate components of extracellular signals, a recent study of the medial temporal area MT showed that LFP activity peaks later than, is partially predicted by, preceding spiking activity, suggesting that modulations of lower frequency LFPs are an epiphenomena of local spiking (Esghaei et al., 2017). However, the extent to which local spiking contributes to LFPs may be brain area- and brain state–dependent (Pesaran et al., 2018), thus the precise nature of the relationship between spikes and LFPs in the PPC during arm position maintenance remains equivocal and a potential focus of future studies.

For the frequency-domain, we expected to see enhanced beta band spectral power at individual sites for only one or both sets of feedback conditions, as was previously observed for spiking activity (VanGilder et al., 2016). This was not generally observed – instead we found that enhanced beta power was ubiquitous across recording sites regardless of sensory condition and final hand location. Moreover, few individual sites showed effects of sensory condition on beta power, though at the population level multisensory suppression of this enhanced beta band activity was observed. What mechanism could account for the prevalence of feedback-independent enhancement of LFP beta power, but *site and feedback-dependent* enhancement of beta power in spiking activity? One possible explanation relates to differences in connectivity patterns of neurons within a given cortical area. Cortical areas receiving strong sensory inputs are thought to contain multiple interconnected subnetworks of neurons that may be selectively responsive to sensory features (Harris and Mrsic-Flogel, 2013). However, all ionic processes (such as transmembrane synaptic inputs) contribute to the extracellular electrical field (Buzsáki et al., 2012; Einevoll et al., 2013). Thus in a given cortical area, the spiking activity of individual neurons may be dependent on their subnetwork-specific responses to selective task conditions, whereas the LFP signal reflects the activity of any of the overlapping subnetworks engaged by a task. Furthermore, spiking activity is dependent on the unique biophysical properties of the neurons within a particular subnetwork, and this may influence which neurons are ultimately entrained by oscillatory synaptic input (Wilson et al., 2018).

One of the overriding principles of MSI, based on numerous behavioral and computation studies, is that that sensory cues are weighed according to their relative reliabilities in a given context (Fetsch et al., 2013). The experiments described here did not systematically alter the reliability of either visual or proprioceptive inputs, thus it is difficult to determine the extent to which the suppression of beta band activity at the population observed here can be attributed to bottom-up, stimulus-driven processes. However, MSI is influenced not only by relative signal reliabilities but also top-down attentional processes (Choi et al., 2018; Keil and Senkowski, 2018; Limanowski and Friston, 2020). Effects of bottom up versus top down processes on MSI have been observed to be frequency dependent, with bottom-up processing being reflected in frequency bands greater than 30 Hz (e.g. gamma band), and top-down processing reflected in lower bands such as beta (Siegel et al., 2012; Keil and Senkowski, 2018). Specific effects such as enhancement or suppression however may also be task and area/network dependent. For example, in an audio-visual congruence task, Friese et al (2016) found that attention led to *increased* gamma band activity but *decreased* beta-band activity in early sensory cortex areas (Friese et al., 2016).

Effects of endogenous attention on MSI have particular relevance to the current results. The data reported here were obtained from recordings of the superficial cortex of the SPL, an area that is believed to be strong proprioceptive inputs from primary somatosensory areas (Cavada and Goldman-Rakic, 1989; Andersen et al., 1990; Caminiti et al., 1996). On U trials, beta band LFP was elevated relative to that on B trials. This could reflect the fact that in order to maintain position under these conditions, animals needed to strongly attend to signals provided by the proprioceptive (and motor) systems. On B trials, where vision was also available for position monitoring, attention to proprioception was likely not as critical. Thus, the reduction in beta activity could reflect an attention-driven shift in the balance of activation among cortical areas involved in position monitoring, from those that are more dominantly proprioceptive to those that are more visual in nature (Limanowski and Friston, 2020).

### Feedback-specific differences in directional/positional tuning

Evoked activity and beta band power were tuned to arm location at many individual sites and at the population level, consistent with evidence that the SPL in primates is involved in the multisensory representation of arm position (Graziano et al., 2000; Lloyd et al., 2003; Pellijeff et al., 2006; Shi et al., 2013; VanGilder et al., 2016) and arm postural control (Lacquaniti et al., 1995; Wolpert et al., 1998; Pellijeff et al., 2006; Parkinson et al., 2010). Critically, however, tuning to hand location often differed between unimodal and bimodal trials. Tuning is not currently believed to be a static feature of neural responses, as tuning curves of individual neurons can change as a function of time, learning, attention, and differences in kinematics/dynamics, among other factors (Donoghue et al., 1998; Li et al., 2001; Paz and Vaadia, 2004; Sergio et al., 2005; Churchland and Shenoy, 2007; Hatsopoulos et al., 2007; Jarosiewicz et al., 2008; Stevenson et al., 2011). In the current experiments, animal’s performed a task on which they were highly trained, thus it is unlikely that learning related factors contributed strongly to condition-dependent tuning differences. In addition, data were analyzed during the same ‘holding’ epoch for both conditions, and neither mean arm position nor position variability differed between conditions. Although it is conceivable that, due to the kinematic redundancy of the arm, different arm configurations were used for the same location in the two conditions, these differences were likely slight, thus it is also unlikely that biomechanical factors contributed strongly to tuning differences between conditions. Lastly, it’s unlikely that the tuning changes observed here were due to temporal factors, as the conditions were run concurrently and were randomly interleaved on a trial by trial basis. This suggests that multisensory interactions altered the tuning properties of some sites (and neurons (Shi et al. 2013)) in the SPL on very short time scales in these experiments.

What could account then for the observed differences in tuning between sensory conditions? As discussed above, attention-driven shifts in the balance of activation among cortical areas could be responsible for the multisensory suppression of beta band activity observed here, thus the notion that tuning differences were also drive by attention is certainly plausible. Another possibility is that differences in tuning could reflect context-dependent encoding of hand position (Buneo and Andersen, 2006; Buneo and Soechting, 2009). That is, in the absence of continuous and reliable visual feedback (e.g., for movements generated in the dark or following long delays without feedback) hand location is determined largely by proprioceptive feedback (and/or forward modeling) and is therefore likely to be encoded in a body-centered frame of reference. In contrast, when the hand is visible, hand location could be remapped from body to eye-centered coordinates, or be encoded in both reference frames simultaneously. Evidence from previous studies focusing on the SPL and dorsal premotor cortex are consistent with the idea that arm movement and posture related variables can be encoded in multiple frames of reference simultaneously(Buneo et al., 2002; Pesaran et al., 2006; Buneo and Andersen, 2012), thus it is conceivable that a similar phenomenon underlies the feedback-specific tuning observed here.

## Acknowledgments

The authors wish to thank Rachele McAndrew for technical assistance.

